# Conservation of symbiotic signalling across 450 million years of plant evolution

**DOI:** 10.1101/2024.01.16.575147

**Authors:** Tatiana Vernié, Mélanie Rich, Tifenn Pellen, Eve Teyssier, Vincent Garrigues, Lucie Chauderon, Lauréna Medioni, Fabian van Beveren, Cyril Libourel, Jean Keller, Camille Girou, Corinne Lefort, Aurélie Le Ru, Didier Reinhardt, Kyoichi Kodama, Syota Shimazaki, Patrice Morel, Junko Kyozuka, Malick Mbengue, Michiel Vandenbussche, Pierre-Marc Delaux

## Abstract

**Highlight:**

- The common symbiotic pathway is activated during arbuscular mycorrhizal symbiosis in *Marchantia paleacea*
- The three core members of the common symbiotic pathway are essential for symbiosis in *Marchantia paleacea*
- The molecular function of the CCaMK/CYCLOPS module is conserved across land plants
- Symbiotic signalling has been conserved in plants for 450 million years

The colonization of land by plants 450 million years ago revolutionized life on Earth^1^. The fossil record^2^ and genetic evidence in extant species^3^ suggest that this transition was facilitated by interactions with symbiotic arbuscular mycorrhizal (AM) fungi^4^. This ancestral symbiosis relied on the biosynthesis of chemicals by the host plant, both as signals^5^ and as nutrients^3^. In angiosperms, a signalling pathway involving the receptor-like kinase SYMRK/DMI2^6,7^, the Calcium and Calmodulin-dependent protein kinase CCaMK/DMI3^8^ and the transcription factor CYCLOPS/IPD3^9,10^ has been described as the common symbiosis pathway (CSP), essential for the establishment of the AM symbiosis and the root-nodule symbiosis^11^. Phylogenetic and comparative phylogenomic analyses indicated an ancient origin of the CSP, present in all extant land plants forming intracellular symbioses^12–15^. Trans-complementation assays of the angiosperm mutants with orthologs from diverse species further indicated the conservation of the molecular function of the CSP across the embryophytes^9,12,14–16^. However, this correlative evidence did not allow testing the ancestral biological function of the CSP. In this study we demonstrate that SYMRK, CCaMK and CYCLOPS are essential for the colonization by AM fungi in bryophytes, indicating that plants have maintained a dedicated signalling pathway to support symbiotic interactions for 450 million years.

## Results and Discussion

### The CYCLOPS-Responsive Element is a marker of CSP induction

In angiosperms, activation of the CSP (Figure 1A) upon symbiont perception leads to the phosphorylation of the transcription factor CYCLOPS by CCaMK, and the transcriptional activation of its direct target genes^17,18^. This direct activation by phosphorylated CYCLOPS is mediated by *cis-* regulatory elements present in the promoter region of the target genes^17,18^. The first of the four different *cis-*regulatory elements bound by phosphorylated CYCLOPS described so far^17–20^ was identified in the promoter of *NIN* from the angiosperm *Lotus japonicus*. A fusion of this element to a GUS reporter (pCYC-RE:GUS) is activated during infection by rhizobial symbionts forming the root-nodule symbiosis in *L. japonicus*^17,21^ and *Medicago truncatula* (Figure S1A). We hypothesized that this reporter could be directly activated by phosphorylated CYCLOPS irrespective of the symbiotic context, and not specifically during the root-nodule symbiosis. *M. truncatula* hairy-roots transformed with the pCYC-RE:GUS reporter were grown in presence or absence of the Arbuscular Mycorrhizal (AM) fungus *Rhizophagus irregularis*, harvested 6 weeks after inoculation and stained for GUS activity. By contrast with the non-inoculated roots that showed only faint and rare staining, the plants inoculated with *R. irregularis* showed consistent and intense GUS induction (Figure 1B and S1B-G). The intense staining colocalized with the presence of the fungal hypheae and arbuscules (Figure 1B). The pCYC-RE:GUS reporter is thus a marker for the activation of the CSP by rhizobial and AM fungi symbionts in *M. truncatula*.

**Figure 1.**
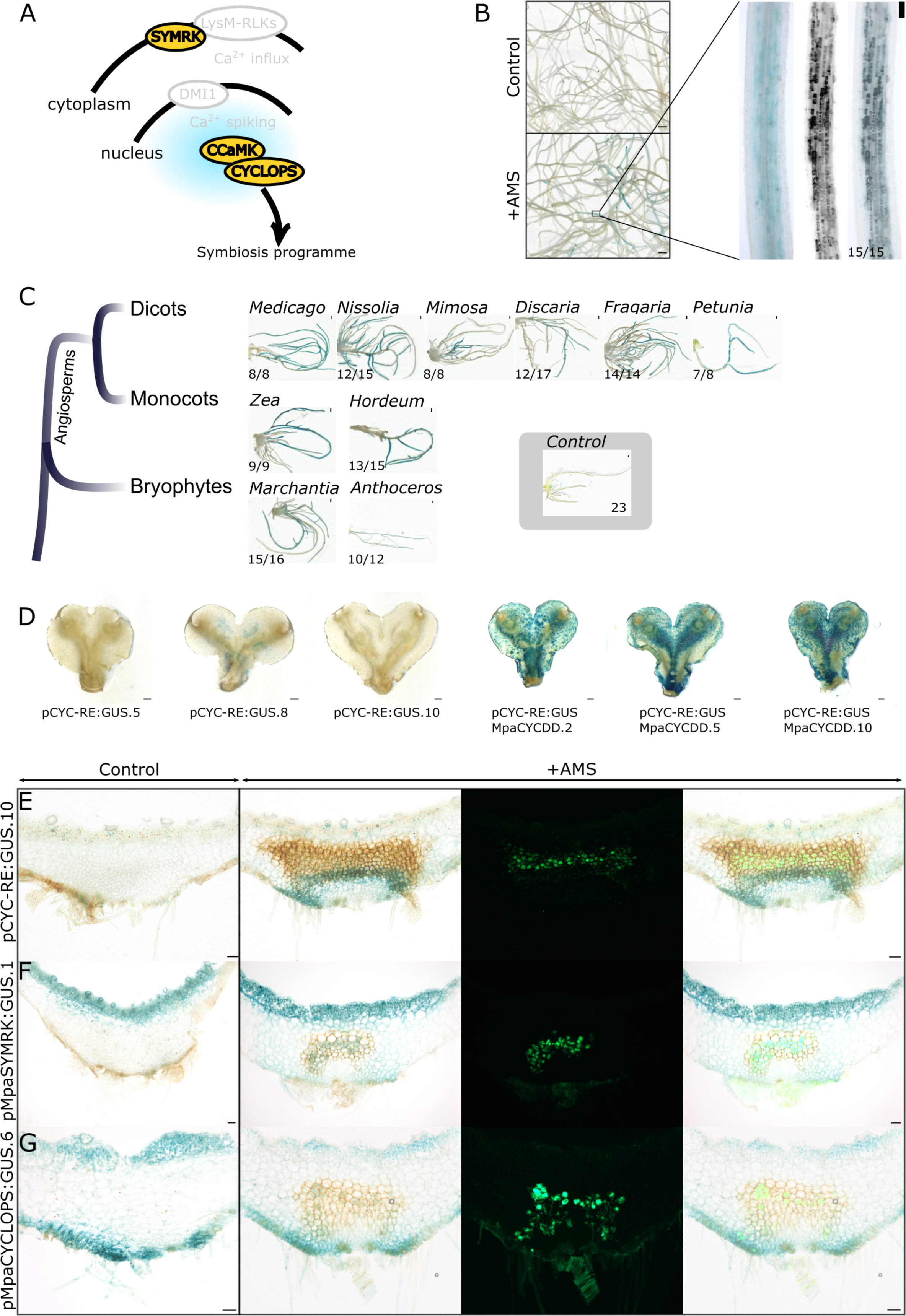
Conservation of pCYC-RE:GUS activation by the CSP transcription factor CYCLOPS in land plants. A. Common Symbiosis Pathway (CSP) in angiosperms. B. pCYC-RE:GUS is activated in response to AMS in *M. truncatula* six weeks post inoculation with *Rhizophagus irregularis*. Whole roots images of GUS-stained inoculated and non inoculated roots are shown (Scale bar 1mm). Faint blue signals could be observed on non inoculated roots whereas intense blue patches were observed on inoculated roots. The zoomed image corresponds to one of these blue patches with a WGA-Alexa Fluor 488 staining revealing AM fungi (black) and an overlay of both images. Scale bar = 100µm. The number of plants with blue signal associated to AM symbiosis is indicated. C. *M. truncatula* roots were transformed with pCYC-RE:GUS and autoactive forms of CYCLOPS (CYCLOPS-DD) orthologs from *M. truncatula*, *Nissolia schottii*, *Mimosa pudica*, *Discaria trinervis*, *Fragaria vesca*, *Petunia axillaris* (*Petaxi*), *Zea mays* (*Zeamay*), *Hordeum vulgare* (*Horvul*), *Marchantia paleacea* (*Marpal*), *Anthoceros agrestis* (*Antagro*) and stained for GUS activity. Control plants correspond to plants transformed only with pCYC-RE:GUS. Plants showing a strong GUS signal out of the total number of observed plants are indicated. For the control, only a faint GUS signal was sometimes observed in the 23 plants observed. Scale bar = 1mm. D. *M. paleacea* transformed lines expressing pCYC-RE:GUS (three independent lines: pCYC-RE:GUS.5, 8 and 10) or pCYC-RE:GUS + MarpalCYCLOPS-DD (three independent lines: pCYC-RE:GUS-MpaCYCDD.2, 5 and 10) are shown after staining for GUS activity. Scale bar = 1mm E. *M. paleacea* transformed with pCYC-RE:GUS (pCYC-RE:GUS.10) without inoculation (Control) or six weeks post inoculation with *R. irregularis*. Thalli were GUS stained and *R. irregularis* is visualized with WGA-Alexa Fluor 488. Bright field, Alexa Fluor 488 and overlay are shown. Scale bar = 100µm. F. *M.* paleacea transformed with pMpaSYMRK:GUS without inoculation (Control) or six weeks post inoculation with *R. irregularis*. Thalli were GUS stained and *R. irregularis* is visualized with WGA-Alexa Fluor 488. Bright field, Alexa Fluor 488 and overlay are shown. Scale bar = 100µm. G. *M.* paleacea transformed with pMpaCYCLOPS:GUS without inoculation (Control) or six weeks post inoculation with *R. irregularis*. Thalli were GUS stained and *R. irregularis* is visualized with WGA-Alexa Fluor 488. Bright field, Alexa Fluor 488 and overlay are shown. Scale bar = 100µm.

To determine whether this activation is specific to the phosphorylation of MtCYCLOPS, we co-expressed in *M. truncatula* hairy roots the pCYC-RE:GUS reporter together with versions of CYCLOPS from various species mimicking the phosphorylation by CCaMK (CYCLOPS-DD^17^). Overexpression of *CYCLOPS-DD* from *M. truncatula, Mimosa pudica* or *Discaria trinervis* which are all able to form both the root-nodule and AM symbioses induced the pCYC-RE:GUS reporter in *M. truncatula* hairy roots (Figure 1C). The same activation was observed when overexpressing *CYCLOPS-DD* from *Fragaria vesca* and *Nissolia schotii* which belong to lineages that lost the ability to form the root-nodule symbiosis, but retained the AM symbiosis^22^. Finally, CYCLOPS-DD from the AM-hosts dicot *Petunia hybrida*, monocots *Zea mays* and *Hordeum vulgare*, and bryophytes *Marchantia paleacea* (thalloid liverworts) and *Anthoceros agrestis* (hornworts), all induced the pCYC-RE:GUS reporter in the absence of AM fungi (Figure 1C). Activation of the pCYC-RE:GUS is thus not limited to phosphomimetic CYCLOPS from species able to form both the root-nodule and the AM symbioses.

To determine whether the activation of the pCYC-RE:GUS was dependent on the genetic background or the symbiotic abilities of the plant species, we expressed *MtCYCLOPS-DD* and the pCYC-RE:GUS reporter in the root of the legume *Nissolia brasiliensis*. The genus *Nissolia* has lost the ability to form the root-nodule symbiosis but retains the AM symbiosis^22^. As for the expression in *M. truncatula,* overexpression of *MtCYCLOPS-DD* resulted in the activation of the pCYC-RE:GUS reporter in *N. brasiliensis* (Figure S1H-I). Finally, overexpression of *MpaCYCLOPS-DD* was conducted in the liverworts *M. paleacea* leading, again, to the activation of the pCYC-RE:GUS reporter (Figure 1D), while control lines only expressing the pCYC-RE:GUS reporter did not show staining (Figure 1D).

Collectively, these data indicate that the pCYC-RE:GUS is a reliable marker for the activation of the CSP, irrespective of the plant species and the type of symbiosis.

### The CSP is activated during symbiosis in Marchantia paleacea

The bryophyte and vascular-plant lineages diverged *ca.* 450 million years ago. Because of this early split during land-plant evolution, identifying conserved features between representatives of these two lineages allows inferring the biology of their most recent common ancestor, a close relative of the first land plants^23^. Among bryophytes, the liverwort *M. paleacea* is able to engage in AM symbiosis^15,24^ and has emerged as an appropriate model to study the conservation of symbiotic processes in land plants. To determine whether the activation of the CSP is conserved across land plants, we first transformed *M. paleacea* with promoter:GUS fusions for the upstream- and downstream-most components of the CSP, namely *SYMRK* and *CYCLOPS*. The lines were inoculated with *R. irregularis* or mock-treated, harvested six weeks later, and stained. In non-inoculated conditions the pMpaSYMRK:GUS and pMpaCYCLOPS:GUS lines displayed background staining in the upper and lower epidermis. Upon inoculation, an additional expression domain was detected for both genes in the cells hosting arbuscules and in the area just below, where intracellular hyphae develop (Figure 1F-G). To directly test the link between AM symbiosis and the CSP, the *M. paleacea* pCYC-RE:GUS reporter lines were inoculated with *R. irregularis*, grown for six weeks and the activation of the reporter tested by GUS staining. While non-inoculated plants showed barely any GUS signal, inoculated plants displayed a robust signal in the part of the thallus hosting intracellular hyphae (Figure 1E). By contrast with *M. truncatula* roots, no signal was observed in the area hosting mature arbuscules, just above the intracellular hypheae (Figure 1E). While in *M. paleacea* the colonization is spatially well-defined, development of the symbiotic structures in *M. truncatula* is asynchronous, mixing cells hosting intracellular hypheae and arbuscules cells. This difference in the zonation of colonization may explain the different pattern observed for the pCYC-RE:GUS reporter. Exploring the induction of this reporter in other host species will allow determining whether this difference is due to lineage specificities or represent two widely distributed patterns.

These results indicate that the CSP is activated during intracellular colonization by AM fungi in *M. paleacea*.

### The symbiotic function of SYMRK is conserved across land plants

The activation of the CSP during AM symbiosis in *M. palacea* is yet another correlative evidence for the conserved symbiotic role of this signalling pathway in land plants. To directly test this role, we generated nine *symrk* mutant alleles in *M. paleacea* using CRISPR/Cas9 (Figure 2A-and S2). Four alleles lead to non-sense mutations coding for predicted truncated proteins (Figure 1B and S2). The other five alleles displayed mis-sense mutations and small deletions that left the downstream original reading frame intact (Figure 1B and S2). The nine mutants were inoculated with *R. irregularis* in parallel with a line transformed with an empty vector (control line). While 96% of the control line thalli were colonized and showed arbuscules five weeks after inoculation, none of the four non-sense alleles lines showed signs of colonization (Figure 2B-C). Among the five mis-sense mutants two were not colonized, and the other three showed a strong quantitative reduction in colonization (Figure 2B). Microscopy confirmed the colonization defect in the *symrk* mutant lines, and the presence of fully developed infection units harbouring arbuscules in the control line (Figure 2C). The consistent AM symbiosis defect in the *M. paleacea symrk* mutant lines is similar to the phenotypes observed in diverse *symrk* mutant of dicots^6,7^ and monocots^25,26^ in which colonization by AM fungi is fully abolished.

**Figure 2.**
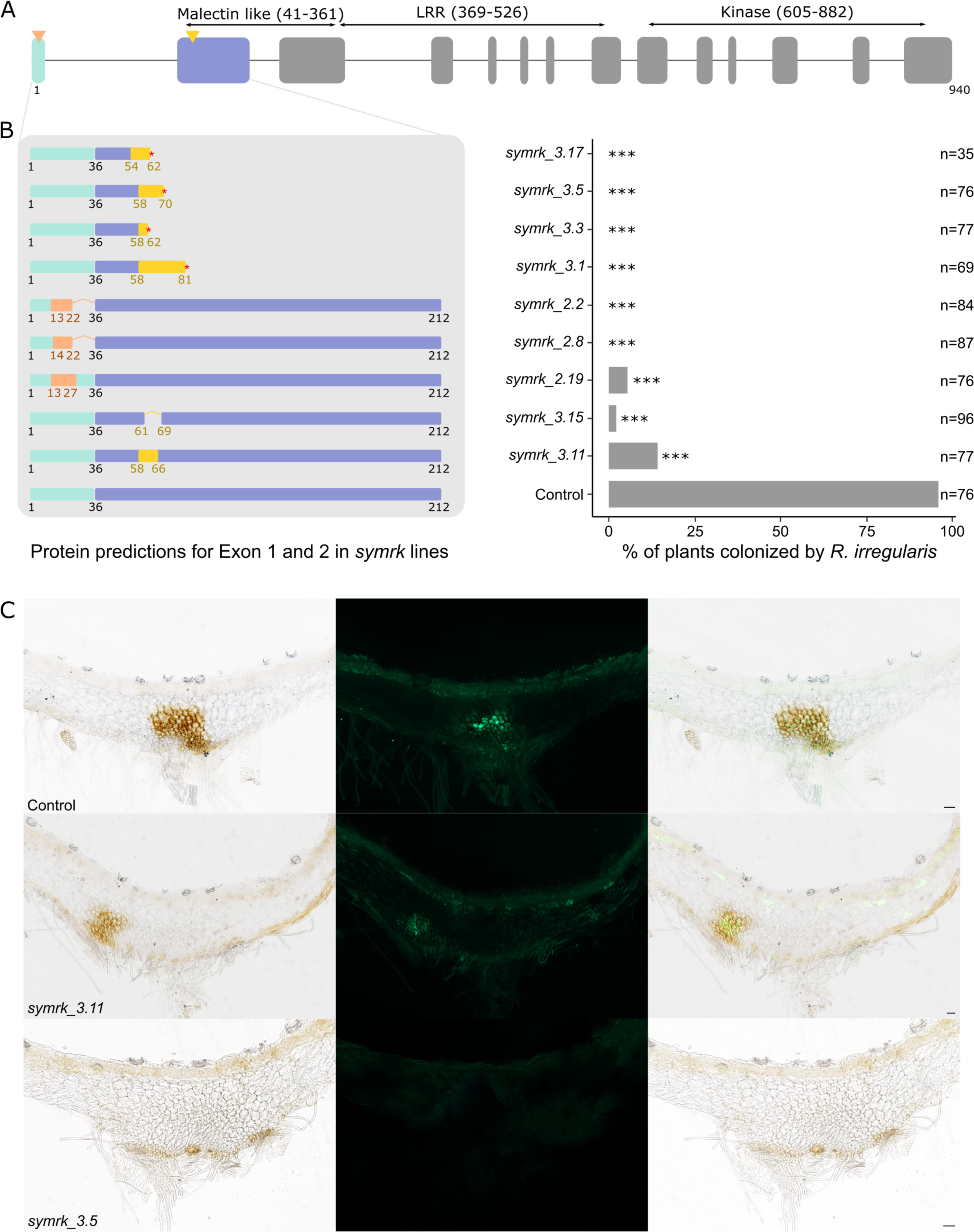
SYMRK is essential fo Arbuscular Mycorrhizal Symbiosis in *Marchantia paleacea.* A. Exons/introns structure of the *MpaSYMRK* genomic sequence. The number of amino acids and predicted domains are indicated. CRISPR/Cas9 was conducted on exon 1 and 2 (arrowheads indicate position of sgRNAs). B. Colonization rates of *M. paleacea symrk* mutant lines at six weeks post inoculation with *R. irregularis*. Predicted protein structures are indicated on the left for the two first exons. Red asterisks indicate the presence of a premature stop codon in the mutant line. *** = statistical difference (p<0,001) calculated with a pairwise comparison of proportions (Chi2) to the control line and a BH p-value adjustment. n= number of observed thalli. C. Transversal sections of *symrk* and control lines six weeks post inoculation with *R. irregularis*. *R. irregularis* is visualized with WGA-Alexa Fluor 488. Bright field, Alexa Fluor 488 and overlay are shown for each line. Scale bar=100µm. *R. irregularis* is observed in control and *symrk_3.11* lines.

Altogether, these data indicate that the biological role of SYMRK for the establishment of the AM symbiosis is conserved across land plants.

### The symbiotic function of the CCaMK/CYCLOPS module is conserved across land plants

Downstream of SYMRK, CCaMK and CYCLOPS act as a module triggering the initial steps of the symbiotic response. In legumes and the monocots rice and barley, CCaMK is essential for AM symbiosis, while *cyclops* mutants display phenotypes ranging from strong reduction in colonization rate to the absence of AM fungi^9,25,27^. Here, we added to the range of tested angiosperms *ccamk* and *cyclops* mutants from a dicot that do not form the root-nodule symbiosis, the Solanaceous species *Petunia hybrida*. Intracellular colonization was neither observed in the *ccamk* mutant nor in the *cyclops* mutant, while the wild-type siblings were well colonized (Table S1) confirming the important role of CCaMK and CYCLOPS for AM symbiosis in angiosperms. Next, we generated thirteen *ccamk* and seven *cyclops* mutants in *M. paleacea* by CRISPR/Cas9 (Figure 3-4 and S3-4). Following inoculation with *R. irregularis*, twelve of the *ccamk* mutants, showed no signs of colonization after five weeks, while the control line (44/54 plants) and a *ccamk* mutant with only a small *in frame* deletion in the sequence preceding the kinase domain (32/40 plants) were normally colonized (Figure 3B). Microscopy confirmed the absence of colonization in the *ccamk* mutant lines, and the presence of fully developed infection units harbouring arbuscules in the control line (Figure 3C). This demonstrates the essential symbiotic role of *ccamk* in *M. paleacea*.

**Figure 3.**
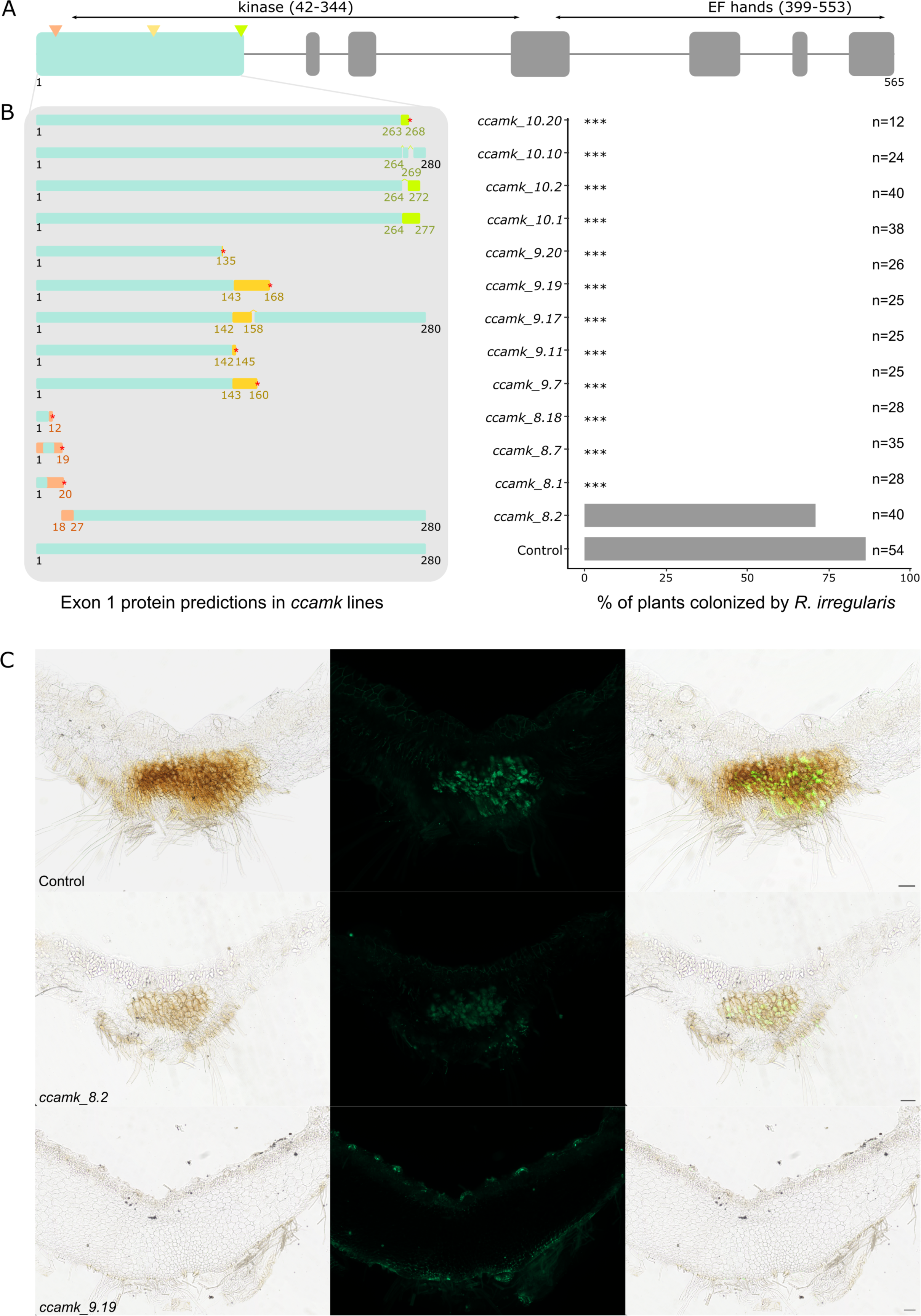
CCaMK is essential for Arbuscular Mycorrhizal Symbiosis in *Marchantia paleacea.* A. Exons/introns structure of the *MpaCCaMK* genomic sequence. The number of amino acids and predicted domains are indicated. CRISPR/Cas9 was conducted on exon 1 (arrowheads indicate position of sgRNAs). B. Colonization rates of *M. paleacea ccamk* lines at six weeks post inoculation with *R. irregularis*. Predicted protein structures are indicated on the left for the first targeted exon. Red asterisks indicate the presence of a premature stop codon in the mutant line. *** = statistical difference (p<0,001) calculated with a pairwise comparison of proportions (Chi2) to the control line and a BH p-value adjustment. n= number of observed thalli. C. Transversal sections of *ccamk* and control lines six weeks post inoculation with *R. irregularis*. *R. irregularis* is visualized with WGA-Alexa Fluor 488. Bright field, Alexa Fluor 488 and overlay are shown for each line. Scale bar=100µm. *R. irregularis* is observed in control and *ccamk_9.19* lines.

**Figure 4.**
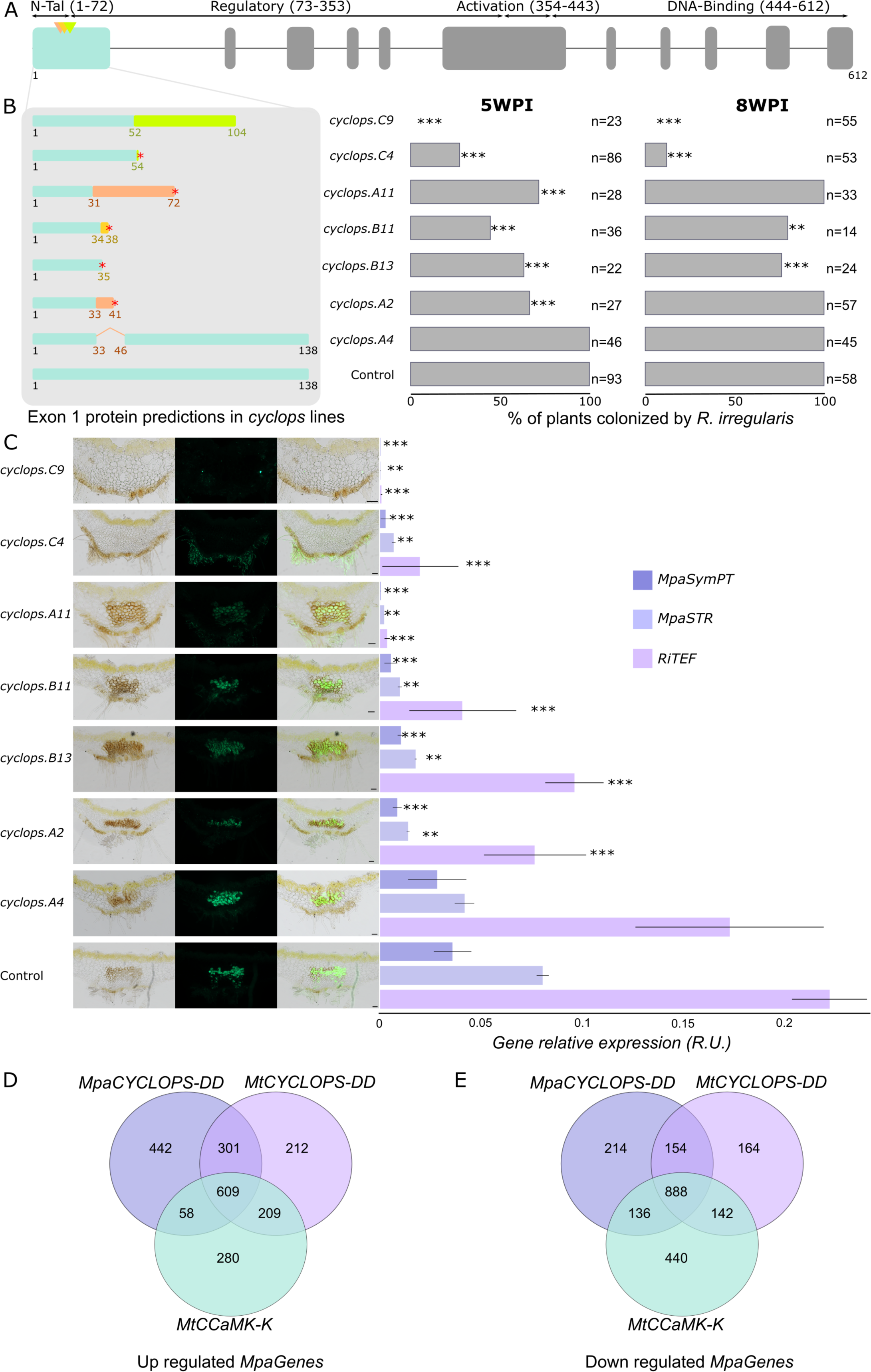
CYCLOPS is important for Arbuscular Mycorrhizal Symbiosis in *Marchantia paleacea.* A. Exons/intron structure of the *MpaCYCLOPS* genomic sequence. Number of amino acids and predicted domains are indicated. CRISPR/Cas9 was conducted on exon 1 (arrowheads indicate position of sgRNAs). B. Colonization rates of *M. paleacea cyclops* lines at five and eight weeks post inoculation with *R. irregularis*. Predicted protein structures are indicated on the left for the first exon. Red asterisks indicate the presence of a premature stop codon in the mutant line. *** and ** indicate statistical difference (p<0,001 and p<0.05 respectively) calculated with a pairwise comparison of proportions (Chi2) to the control line and a BH p-value adjustment. n= number of observed thalli. C. Transversal sections of *M. paleacea cyclops* and control lines ight weeks post inoculation with *R. irregularis*. *R. irregularis* is visualized with WGA-Alexa Fluor 488. Bright field, Alexa Fluor 488 and overlay are shown for each line. Scale bar=100µm. *R. irregularis* is observed inside all thalli of all lines except *cyclops.C9* and *cyclops.C4*. On the right panel, the bars correspond to the expression levels of *MpaSymPT*, *MpaSTR* and *RiTEF* analyzed by qRT–PCR on thalli five weeks post inoculation with *R. irregularis*. Data were normalized to *M. paleacea* housekeeping gene MpaEF1. Error bars represent SE (*n* = 3). Asterisks indicate statistically significant differences (Student’s *t-*test: ∗ *p-value* ≤ 0.1, ∗∗ *p-value* ≤ 0.05, and ∗∗∗ *p-value* ≤ 0.001) compared with the control lines. D. Venn diagrams of up-regulated genes in *M. paleacea* overexpressing *MpaCYCLOPS-DD*, *MtCYCLOPS-DD*, or *MtCCaMK-Kin*, respectively (FDR≤0.05). E. Venn diagrams of down regulated genes in *M. paleacea* overexpressing *MpaCYCLOPS-DD*, *MtCYCLOPS-DD*, or *MtCCaMK-Kin*, respectively (FDR≤0.05).

Five weeks after inoculation with *R. irregularis*, the phenotypes of the *cyclops* frameshift mutants ranged from moderately to strongly reduced colonization, or to a total lack of colonization (Figure 4B). One allele, only affected by a 12nt in-frame deletion, was colonized to a similar level than the control line (Figure 4B). Similar phenotypes were still observed eight weeks after inoculation (Figure 3B). The observed difference in the strength of the phenotypes between *cyclops* mutants correlates with the position of the CRISPR/Cas9-induced mutations (Figure 4 and S4). While the alleles showing lack of, or strongly reduced, colonization were mutated in a domain conserved across all embryophytes (Figure 4A-4B and S4-5), alleles with reduced colonization were mutated closer to the N-terminal part of the protein, in a domain conserved across bryophytes, ferns and gymnosperms, but missing from the angiosperms (Figure S5). This additional domain may thus have a non-essential function. Presence of alternative start codons downstream the mutation present in the weak alleles (Figure S5) supports this hypothesis, although further experiments are required to test it. Strongly reduced colonization was also observed in two additional frameshift mutant lines generated in *M. paleacea ssp diptera* (Figure S6). To consolidate the quantification of the phenotypes, RNA was extracted from the seven *M. paleacea cyclops* mutants and the empty vector control line five weeks after inoculation with *R. irregularis*, and the expression of the AM-responsive phosphate (*MpaSymPT*) and lipid (*MpaSTR*) transporters was monitored by qRT-PCR. Expression of the *R. irregularis* housekeeping gene *RiTEF* was quantified as a proxy for fungal abundance. The expression level of *SymPT*, *STR* and *RiTEF* mirrored the observed colonization rates, confirming that the functionality of the AM symbiosis is impaired in the *cyclops* frameshift mutants (Figure 4C).

The symbiotic defects observed in *M. paleacea ccamk* and *cyclops* mutants, reminiscent of the ones observed in angiosperms, indicate that the biological function of these two genes as regulators of the AM symbiosis is conserved across land plants.

### The molecular function of the CYCLOPS/CCaMK module is conserved across land plants

Complementation of the symbiotic defects of angiosperms *ccamk* and *cyclops* mutants by their bryophyte orthologs supports the conservation of the molecular function of this module across plant lineages^12,14,15^. In angiosperms, CCaMK phosphorylates CYCLOPS following symbiont-induced nuclear calcium spiking^9^. In both *M. paleacea* and angiosperms expression of a gain of function version of *M. paleacea* CYCLOPS mimicking this phosphorylation (*MpaCYCLOPS-DD*) leads to the activation of the pCYC-RE:GUS reporter (Figure 1C-D). If the CYCLOPS/CCaMK module is indeed conserved across plant lineages, we reasoned that overexpression of *MpaCYCLOPS-DD*, *MtCYCLOPS-DD and MtCCaMK-Kin,* an autoactive version of CCaMK^15,28^, should result in similar transcriptomic signatures relative to control lines. To test this, *M. paleacea* lines overexpressing either of these three constructs were generated, their transcriptome determined by RNAseq, and compared to lines transformed with an empty vector to identify differentially expressed genes (Figure 4, Table S2). All three constructs lead to very significant transcriptomic changes, ranging from 1156 to 1410 up-regulated genes, and 1348 to 1606 down-regulated genes (Figure 4D-4E, Table S2). Among the genes up-regulated in response to *MpaCYCLOPS-DD,* 910 and 667 were also found up-regulated in response to *MtCYCLOPS-DD* and *MtCCaMK-K* respectively (Figure 4D). A similar trend was observed for the down-regulated genes (1042 and 1024 respectively, Figure 4E). These overlaps were significantly higher than expected by chance (Table S3).

Mirroring the trans-complementation of angiosperm mutants with liverwort sequences^12,14,15^, these results indicate that, similar to the conservation of the biological role of the CYCLOPS/CCaMK module, the molecular function of these two component of the CSP is conserved across land plants.

## Conclusion

As described over the last two decades for angiosperms, the CSP components *SYMRK*, *CCaMK* and *CYCLOPS* are essential for AM symbiosis in the liverwort *M. paleacea*. To explain this conserved role, the most parsimonious hypothesis suggests that the most recent common ancestor of the angiosperms and the bryophytes already used the CSP to engage and associate with AM fungi. In other words, our data indicate that plants have relied on a conserved symbiotic signalling pathway for 450 million years. Phylogenomic data indicate that this pathway may subsequently have been co-opted for other intracellular symbioses with diverse fungi and nitrogen-fixing bacteria^15^. Genetics in legumes support this hypothesis for the root-nodule symbiosis^11^. How transitions from one symbiotic type to another occurred by co-opting the very ancient CSP represents the next challenge to be deciphered. Such an understanding may open the possibility to expand the symbiotic abilities of crops species by mimicking evolution.

## Acknowledgements

We thank the genotoul bioinformatics platform Toulouse Occitanie (Bioinfo Genotoul, https://doi.org/10.15454/1.5572369328961167E12) for providing computing resources. We acknowledge the TRI-FRAIB imaging facility, member of the national infrastructure France-BioImaging supported by the French National Research Agency (ANR-10-INBS-04). Research performed at LRSV was also supported by the ‘Laboratoires d’Excellence (LABEX)’ TULIP (ANR-10-LABX-41) and the ‘École Universitaire de Recherche (EUR)’ TULIP-GS (ANR-18-EURE-0019). L.M., C.G., T.V., J.K., C.L. and P-M.D were supported by the project Engineering Nitrogen Symbiosis for Africa (ENSA) funded through a grant to the University of Cambridge by the Bill and Melinda Gates Foundation (OPP1172165) and the UK Foreign, Commonwealth and Development Office as Engineering Nitrogen Symbiosis for Africa (OPP1172165). This project received funding from the European Research Council (ERC) under the European Union’s Horizon 2020 research and innovation programme (grant agreement no. 101001675 - ORIGINS) to P.-M.D.

## Author contributions

T.V., P-M.D, M.K.R., M.V., J.Ky., M.M., D.R. designed and coordinated the experiments. T.V., A.L-R, P.M., M.V., F.VB., M.K.R., E.T., J.Ke., C.Le., C.Li., C.G., T.P., L.C., V.C., K.K. and S.S. conducted experiments. P-M.D. and T.V. wrote the manuscript. P-M.D. coordinated the project.

## Data and resource availability

### Lead contact

Requests for resources and further information should be directed towards Pierre-Marc Delaux.

## Materials availability

Plasmids and transgenic lines generated in this study are available. For the transfer of transgenic material, appropriate information on import permits will be required from the receiver.

## Data and code availability

RNAseq reads were deposited on the SRA with the Bioproject number PRJNA1051818. This paper does not report original code. Any additional information required to reanalyze the data reported in this paper is available from the lead contact upon request.

## Material and methods

### Phylogeny

To reconstruct the phylogeny of CYCLOPS, we recovered protein sequences from a variety of land plants using hmmscan from HMMER v3.4^29^ with the HMM profile of the CYCLOPS domain (IPR040036). A set of 37 protein sequences was aligned using MAFFT v7.520 with the E-INS-i method^30^. The phylogeny was reconstructed using IQ-TREE v2.2.2.3 with the model LG+C20+F+G and support was provided with 1,000 ultrafast bootstrap replicates^31–34^. The tree was rooted between bryophytes and vascular plants, and the tree and alignment were visualized using ETE3 v3.1.2^35^.

### Cloning

The Golden Gate modular cloning system^36,37^ was used to prepare the plasmids as described in Rich et al. ^3^for all constructs, except for pMpaSYMRK:GUS. Levels 0, 1 and 2 used in this study are listed in Table S4 and held for distribution in the ENSA project core collection (https://www.ensa.ac.uk/). Sequences were domesticated (listed in Table S4), synthesized and cloned into pMS (GeneArt, Thermo Fisher Scientific, Waltham, USA).

Gateway system was used to construct pMpaSYMRK:GUS. pMpaSYMRK (2.4kb) was amplified with GGGGACAAGTTTGTACAAAAAAGCAGGCTTCGCTTCTCAGAAACAACTCTA and GGGGAGCCACTTTGTACAAGAAAGCTGGGTCGTTCTGCTTCAAACCGAGAC and cloned by BP in pDOn207 and then into pMDC164 by LR^38^.

#### Generation of CRISPR mutants in M. paleacea ssp. Paleacea

Constructs containing the *Arabidopsis thaliana* codon optimized Cas9^39^ under the MpoEF1a promoter and two guide RNA under the *M. paleacea* or *M. polymorpha* U6 promoter were transformed in *M. paleacea*. A total of nine alleles of *symrk* (*Marpal_utg000051g0090241*), thirteen of *ccamk* (*Marpal_utg000137g0173321*) and seven of *cyclops* (*Marpal_utg000051g0091871*) were genotyped and selected for phenotyping (Table S5 and S6).

#### Generation of CRISPR mutants in M. paleacea ssp. Diptera

To generate mutants of CYCLOPS in *Marchantia paleacea ssp. diptera* (*cyclops.D1*, *cyclops.D2*), plants were transformed with the construct containing Arabidopsis-codon-optimized Cas9 fused with MpoEF1a promoter and a guide RNA (GCTCGAACCATATTCATG) fused to the MpoU6-1 promoter. Two edited lines were selected for phenotyping.

### Medicago assays

Constructs were transformed in *Agrobacterium rhizogenes* A4TC24 by electroporation. Transformed strains were grown at 28°C in Luria-Bertani medium supplemented with rifampicin and kanamycin (25 µg/mL). *M. truncatula* Jemalong A17 roots, were transformed with the different CYCLOPS-DD orthologs and the pCYC-RE:GUS (Table S4) as described by Boisson-Dernier et al.^40^, and grown on Fahraeus medium for 2 months, selected with the DsRed marker present in all the constructs and GUS stained as in Vernié *et al.* 2015^41^.

For nodulation and mycorrhization assays, *M. truncatula* plants with DsRed-positive roots were transferred to pots containing Zeolite substrate (50% fraction 1.0-2.5mm, 50% fraction 0.5-1.0-mm, Symbiom). For nodulation assays, plants were watered with liquid Fahraeus medium. Wild-type *S. meliloti* RCR2011 pXLGD4 (GMI6526) was grown at 28°C in tryptone yeast medium supplemented with 6 mM calcium chloride and 10 µg/mL tetracycline, rinsed with water and diluted at OD600=0.02. Each pot was inoculated with 10 ml of bacterial suspension. For mycorrhization assays, each pot was inoculated with *ca*. 500 sterile spores of *Rhizophagus irregularis* DAOM 197198 (Agronutrition, Labège, France) and grown with a 16 h/8 h photoperiod at 22 °C/20 °C. Pots were watered once per week with Long Ashton medium containing 15 μM phosphate^42^.

### Nissolia assays

*Nissolia brasiliensis* seeds provided by CIAT (Programade Recursos Geneticos, Valle, Colombia) were scarified with sulfuric acid for 5min and surface-sterilized with bleach for 1min. Seeds were washed 5 times with H_2_O at each step. Seeds were placed onto 0.8% (w/v) agar plates in a growth chamber (25°C) under dark conditions for 3 days. Germinated seedlings were pierced with a needle that had been previously dipped in the *A. rhizogenes* inoculum at OD=0.03, and placed on Fahraeus medium plates in a 25°C growth chamber (16h light/8h dark). After 10 days, the untransformed roots (DSred-negative) were removed with a scalpel blade. After one month, transformed roots were screened for DsRed, and GUS stained as indicated for *M. truncatula*.

### Petunia assays

#### Petunia genotyping

Petunia LY3784 (*cyclops-1*) and 86-5 (*ccamk-1*) mutants were identified by searching a sequence-indexed dTph1 transposon database^43^. Exact insert positions (expressed in base pairs downstream of the ATG start codon with the coding sequence as a reference) were determined by aligning the dTph1 flanking sequences with the genomic and cDNA sequences. All *in silico*-identified candidate insertions were confirmed by PCR-based genotyping of the progeny from the selected insertion lines, using primers flanking the dTph1 transposon insertions (ATGCAGCATAATATACCAGGAAATG and TGGGCTGGTTAGTAGTTTCAC for *CYCLOPS*, AAATTTTCCACACTCTTGATCAAACTC and AGCCACCTCTTCCAAGTATGTC for *CCaMK*). The following thermal profile was used for segregation analysis PCR: 10 cycles (94°C for 15 s, 68°C for 20 s minus 1°C/cycle, 72°C for 30 s), followed by 40 cycles (94°C for 15 s, 58°C for 20 s, and 72°C for 30 s). The different insertion mutants were further systematically genotyped in subsequent crosses and segregation analyses. PCR products were analyzed by agarose gel electrophoresis. *cyclops-1* has an insertion at 609bp, *ccamk-1* at 262bp.

#### Mycorrhization tests in Petunia hybrida

Seeds were germinated by sowing in pots with wet soil, at the surface (without covering). Then, a mini-greenhouse was placed over the pots to keep a high humidity and seeds were left to germinate in a growth chamber (25°C day/22°C night, 60% humidity, 16h/8h day/night). Germinated seedlings were transferred to zeolite (50% fraction 1.0-2.5mm, 50% fraction 0.5-1.0-mm, Symbiom) soaked in Long-Ashton solution containing 15 μM of phosphate and inoculated with *ca.* 500 spores/pot of *R. irregularis* DAOM 197198 (Agronutrition, Labège, France). Plants were grown in a growth chamber (25°C day/22°C night, 60% humidity, 16h/8h day/night) and watered regularly with the Long-Ashton solution. Root systems were harvested after 5 weeks and stained with ink. Mycorrhization was quantified using the grid intersection method^44^.

### Marchantia assays

#### Marchantia paleacea ssp paleacea transformation

Gemmae collected from axenic *M. paleacea* were grown in ½ strength Gamborg B5 media (G5768, Sigma) pH 5.7, 1.4% bacteriological agar (1330, Euromedex) for 4-5 weeks.

For each construct, 15-25 gemmalings were blended for 15 seconds in a sterile, 250ml stainless steel, bowl (Waring, USA) in 10 ml of 0M51C medium (KNO_3_ 2g/L, NH_4_NO_3_ 0.4g/L, MgSO_4_ 7H2O 0.37g/L, CaCl_2_ 2H_2_O 0.3 g/L, KH_2_PO_4_ 0.275 g/L, L-glutamine 0.3 g/L, casamino-acids 1 g/L, Na_2_MoO_4_ 2H_2_O 0.25 mg/L, CuSO_4_ 5H2O 0.025 mg/L, CoCl_2_ 6H2O 0.025 mg/L, ZnSO_4_ 7H_2_O 2 mg/L, MnSO_4_ H_2_O 10 mg/L, H_3_BO_3_ 3 mg/L, KI 0.75 mg/L, EDTA ferric sodium 36.7 mg/L, myo-inositol 100 mg/L, nicotinic acid 1 mg/L, pyridoxine HCL 1 mg/L, thiamine HCL 10 mg/L). The blended plant tissues were transferred to 250ml erlenmeyers containing 15 ml of 0M51C and kept at 20°C, 16h light/8h dark, on a shaking table (200 RPM) for 3 days. Co-culture was initiated by adding 100 µL of saturated *Agrobacterium tumefasciens* GV3101 liquid culture and acetosyringone (100 µM final). After 3 days, the plant fragments were washed by decantation three times with water, and plated on ½ Gamborg containing 200 mg/L amoxycilin (Levmentin, Laboratoires Delbert, FR) and 10 mg/L Hygromycin (Duchefa Biochimie, FR).

#### Marchantia paleacea ssp diptera transformation

Transformation was done as in Kodama et al 2022^5^. Parental lines used to generate cyclops mutant were used as controls (Control.F).

#### GUS-staining

Plants, either mock-treated or inoculated with *R. irregularis* for 6 weeks, were harvested. For staining, the GUS buffer is composed of: phosphate buffer (0.1 M), EDTA (5 mM), K_3_Fe(CN)_6_ (0.5 mM), K_4_Fe(CN)6 (0.5 mM), X-Glu (0.25 mg/ml) and H_2_O. After covering the plants with the GUS buffer, the tissues were incubated under vacuum for 5 min (twice), before incubating at 37°C for 12-15h. Several washes were performed with 70% ethanol to remove chlorophyll and clear the tissue. Tissues were stored in an aqueous solution containing EDTA (0.5 M).

#### Mycorrhization tests in Marchantia paleacea

Thalli of *Marchantia paleacea ssp paleacea* and *Marchantia paleacea ssp diptera* were grown on a zeolite substrate (50% fraction 1.0-2.5mm, 50% fraction 0.5-1.0-mm, Symbiom) in 7×7×8 cm pots (five thalli per pot). Each pot was inoculated with *ca*. 1,000 sterile spores of *Rhizophagus irregularis* DAOM 197198 (Agronutrition, Labège, France) and grown with a 16h/8h photoperiod at 22°C/20°C. Pots were watered once a week with Long-Ashton medium containing 15 μM of phosphate.

Six weeks post-inoculation, thalli were cleared of chlorophyll using ethanol 100% for 24 hours, then stored in an aqueous solution containing EDTA (0.5 mM). Cleared thalli were scanned and the presence of the black/purple pigment indicative of colonization scored. Colonization or lack of colonization was confirmed by staining as presented below. Mycorrhization assays were run independently 2-5 times.

#### Microscopy on M. paleacea

Thalli were embedded in 6% agarose and 100µm transversal and horizontal sections were prepared using a Leica vt1000s vibratome. Sections were incubated two days in 10% KOH at 4°C followed by water washes. The sections were then incubated in the staining solution, PBS with 1 µg/ml WGA-Alexa Fluor 488 (Invitrogen) overnight at 4°C. Pictures were taken with a Nikon Eclipse Ti with the camera lens 10x/0.3 and with a Zeiss Axiozoom V16 microscope. Images were processed with ImageJ.

#### qRT-PCR

RNAs of *M. paleacea cyclops* mutant lines or empty-vector control plants were extracted using a Direct-zol RNA MiniPrep Zymo kit according to the supplier’s recommendation on ∼100 mg of ground frozen thalli.

Reverse transcription was performed using M-MLV (Promega, USA) on 500 ng of RNA and qPCR was performed on 5x diluted cDNA in a BioRad CFX opus 384 thermocycler with SYBR Green (Sigma). Relative expression values were calculated using the reference gene *MpaEF1*.

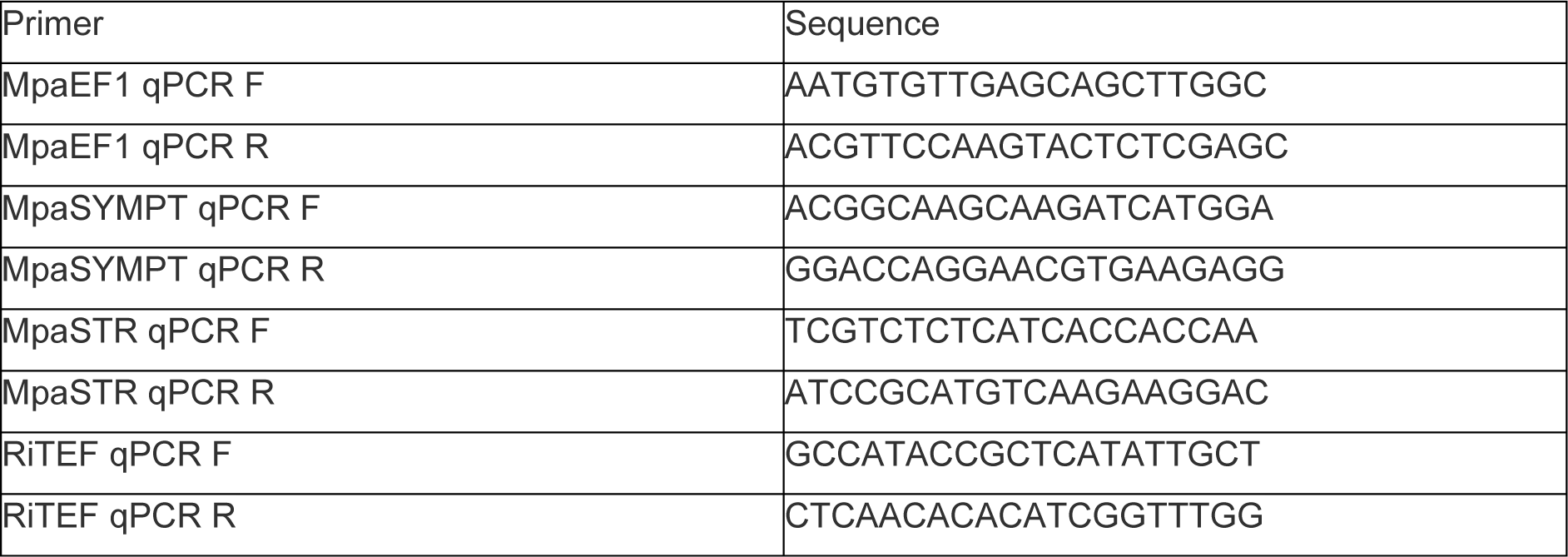

### RNAseq

#### Library preparation

Three independent lines expressing MtCYCLOPS-DD, MtCCaMK-K, MpaCYCLOPS-DD or transformed with an empty vector control (Line 132) were harvested five weeks after transfer to zeolite substrate (50% fraction 1.0-2.5mm, 50% fraction 0.5-1.0-mm, Symbiom) in 7×7×8 cm pots (five thalli per pot). Thali from each pot were pooled in a single sample, flash-frozen and stored at −70°C until extraction. TRI-reagent (Sigma) extraction was performed according to the supplier’s recommendation on ∼100mg of ground frozen thalli. Around 2µg of RNA was treated with RQ1 DNase (Promega, USA) and sent for sequencing to Genewiz/Azenta (Leipzig, Germany). Illumina libraries were prepared with the NEBnext ultra II RNA directional kit and sequenced on a NovaSeq platform.

#### Differential gene expression analysis

All sequenced RNAseq libraries were mapped against the reference genome of *M. paleacea*^3^ using nextflow^45^ (v21.04.1, build 5556) run nf-core/rnaseq^46^ (v3.4, 10.5281/zenodo.1400710) using *-profile debug, genotoul--remove_ribo_rna--skip_qc--aligner star_salmon* options. The workflow used bedtools^47^ (v2.30.0), bioconductor-summarizedexperiment (v1.20.0), bioconductor-tximeta (v1.8.0), gffread^48^ (v0.12.1), picard (v2.25.7), salmon^49^ (v1.5.2), samtools^50^ (v1.13), star^51^ (v2.6.1d), stringtie^52^ (v2.1.7), Trimgalore (v0.6.7, <u>GitHub - FelixKrueger/TrimGalore: A wrapper around Cutadapt and</u> <u>FastQC to consistently apply adapter and quality trimming to FastQ files, with extra functionality for</u> <u>RRBS data</u>), cutadapt^53^ (v3.4) and ucsc (v377). Differencially expressed genes (DEGs) for the different lines were estimated using ‘*edgeR*’^54^ in R (v4.1.2, R Core Team 2021). Briefly, low expressed genes with less than 10 reads across each class of samples were removed. Then, gene counts were normalized by library size and trimmed mean of M-values (*i.e.* TMM) normalization method^55^. We estimated differentially expressed genes (DEGs) by comparing each transformed genotype (MtCYCLOPS-DD, MtCCaMK-K and MpaCYCLOPS-DD) to empty vector plants. Genes were considered differentially expressed when the FDR was below 0.05 (Benjamini-Hochberg correction).

#### Statistical analyses

To estimate if the observed overlap between genes deregulated by the overexpression of the different constructs in *M. paleacea* (MtCYCLOPS-DD, MtCCaMK-K and MpaCYCLOPS-DD) differed from random expectations, we randomly sampled (10,000 times) the same number of genes than the number of genes deregulated in each treatment, and cross-referenced these random datasets to estimate the random overlap. Quantiles metrics were computed for each comparison.

## Supplemental Information

**Supplemental Figure 1. pCYC-RE expression in *M. truncatula* roots**

**A.** The pCYC-RE:GUS signal in *M. truncatula* roots inoculated with *S. meliloti* is localized in young dividing cells (triangle) at the origin of nodule primordium, in young nodule (star) and at the base of developing and 12 days-old nodules (arrow). **B-G.** *M. truncatula* roots expressing pCYC-RE:GUS six weeks post inoculation with *R. irregularis* (B, D, F) or non-inoculated (C, E, G). In inoculated roots the blue signal is stronger and associated with colonized cells, whereas in non-inoculated roots, only some weak and diffuse signal can be observed in a few roots. In F, an overlay with WGA-Alexa Fluor 488 signal associated with *R. irregularis* is shown. Scale bars correspond to 1 cm in B and C, and to 100µm in other panels. **G-H.** *Nissolia brasiliensis* roots transformed with pCYC-RE:GUS (H) and pCYC-RE:GUS + MtCYCLOPS DD (I). Number of plants showing a similar expression pattern out of the total number of transformed plants are indicated.

**Supplemental Figure 2: Genomic and proteic alignments of MpaSYMRK in wild type and *M. paleacea* mutant lines.**

**A** and **B.** Genomic alignment of the first and second exon of *MpaSYMRK*. Pairs of sgRNAs used for CRISPR/Cas9 are underlined in different colors. Premature stop codons are underlined in black.

**A. C.** Proteic alignment of the two first exons of *MpaSYMRK* from the different mutant lines. Alignments were performed using Clustal Omega and visualized in AliView.

**Supplemental Figure 3: Genomic and proteic alignments of MpaCCaMK in wild type and *M. paleacea* mutant lines.**

**A.** Genomic alignment of the first exon of *MpaCCaMK*. Pairs of sgRNAs used for CRISPR/Cas9 are underlined in different colors. Premature stop codons are underlined in black.

**B.** Proteic alignment of the first exon of MpaCCaMK from the different mutant lines. Alignments were performed using Clustal Omega and visualized in AliView.

**Supplemental Figure 4: Genomic and proteic alignments of MpaCYCLOPS in wild type and *M. paleacea ssp paleacea* mutant lines.**

**A.** Genomic alignment of the first exon of *MpaCYCLOPS*. Pairs of sgRNAs used for CRISPR/Cas9 are underlined in different colors. Premature stop codons are underlined in black and the predicted new in-frame start codons are underlined in blue. The N-terminal domain absent from angiosperms is indicated in green, the regulatory domain in black.

**B.** Proteic alignment of MpaCYCLOPS first exon until premature stop codon if a stop codon is present.

**C.** Proteic alignment of MpaCYCLOPS first exon from sequences with new predicted in-frame start codons in the different mutant lines. Alignments were performed using Clustal Omega and visualized in AliView.

**Supplemental Figure 5: Loss of a N-terminal domain in angiosperms CYCLOPS proteins**

CYCLOPS phylogeny reconstructed using maximum likelihood. The untrimmed alignment of the proteins is shown next to the phylogeny. The N-terminal domain is absent (or highly divergent) in angiosperms and is highlighted in green. The regulatory, activation, and DNA-binding domains are highlighted in red, yellow, and blue, respectively. The two phosphorylation sites are shown with a dark blue rectangle; they were both mutated to D in CYCLOPS-DD overexpressing lines. Ultrafast bootstrap support (%; out of 1,000 replicates) is shown at the end of the branches. The scale represents the estimated number of substitutions per site. Sequences that differ from the original annotation are shown with an asterisk (*), and experimentally validated proteins are marked with a yellow star.

**Supplemental Figure 6: *Marchantia paleacea ssp diptera cyclops* lines are impaired for arbuscular mycorrhization with *R. irregularis***

**A.** Proteic alignments of MpaCYCLOPS first exon in wild type and *M. paleacea ssp paleacea* and *ssp diptera* mutant lines. The N-terminal domain absent from angiosperms is indicated in green, the regulatory domain in black. Alignments were performed using Clustal Omega and visualized in AliView.

**B.** The percentage of plants colonized by *R. irregularis* five weeks post inoculation with *R. irregularis* is indicated. *** indicates statistical difference (p-value<0,001) calculated with a pairwise comparison of proportions (Chi2) to the control line (Control.F) and a BH p-value adjustment. n= number of observed thalli.

**C.** Sections of *cyclops* and control lines five weeks post inoculation with *R. irregularis*. *R. irregularis* is visualized with blue ink. Scale bar=100µm.

**Table S1. Number of plants showing root mycorrhizal arbuscules in *petunia* lines *ccamk*, *cyclops* and wild type**

**Table S2. List of differentially regulated genes in response to CYCLOPS-DD and CCaMK-K overexpression in *Marchantia paleacea***

**Table S3. Statistical test of the overlap between CYCLOPS-DD and CCaMK-K-induced transcriptomic changes**

**Table S4. List of constructs used in this study.**

**Table S5. List of primers for CRISPR lines**

**Table S6. *Marchantia paleacea ssp paleacea* selected lines in this study**

