## Supplemental Figures 1 to 6 for "Conservation of symbiotic signalling across 450 million years of plant evolution"

**Supplementary Figure 1: CYC RE expression in *M. truncatula* roots**

**A.** The pCYC-RE:GUS signal in *M. truncatula* roots inoculated with *S. meliloti* is localized in young dividing cells (triangle) at the origin of nodule primordium, in young nodule (star) and at the base of developing and 12 days-old nodules (arrow). **B-G.** *M. truncatula* roots expressing pCYC-RE:GUS six weeks post inoculation with *R. irregularis* (**B, D, F**) or non-inoculated (**C, E, G**). In inoculated roots the blue signal is stronger and associated with colonized cells, whereas in non inoculated roots, only some weak and diffuse signal can be observed in a few roots. In **F**, an overlay with WGA-Alexa Fluor 488 signal associated with *R. irregularis* is shown. Scale bars correspond to 1 cm in **B** and **C**, and to 100µm in other panels. **G-H.** *Nissolia brasiliensis* roots transformed with pCYC-RE:GUS (**H**) and pCYC-RE:GUS + MtCYCLOPS DD (**I**). Number of plants showing a similar expression pattern out of the total number of transformed plants are indicated.

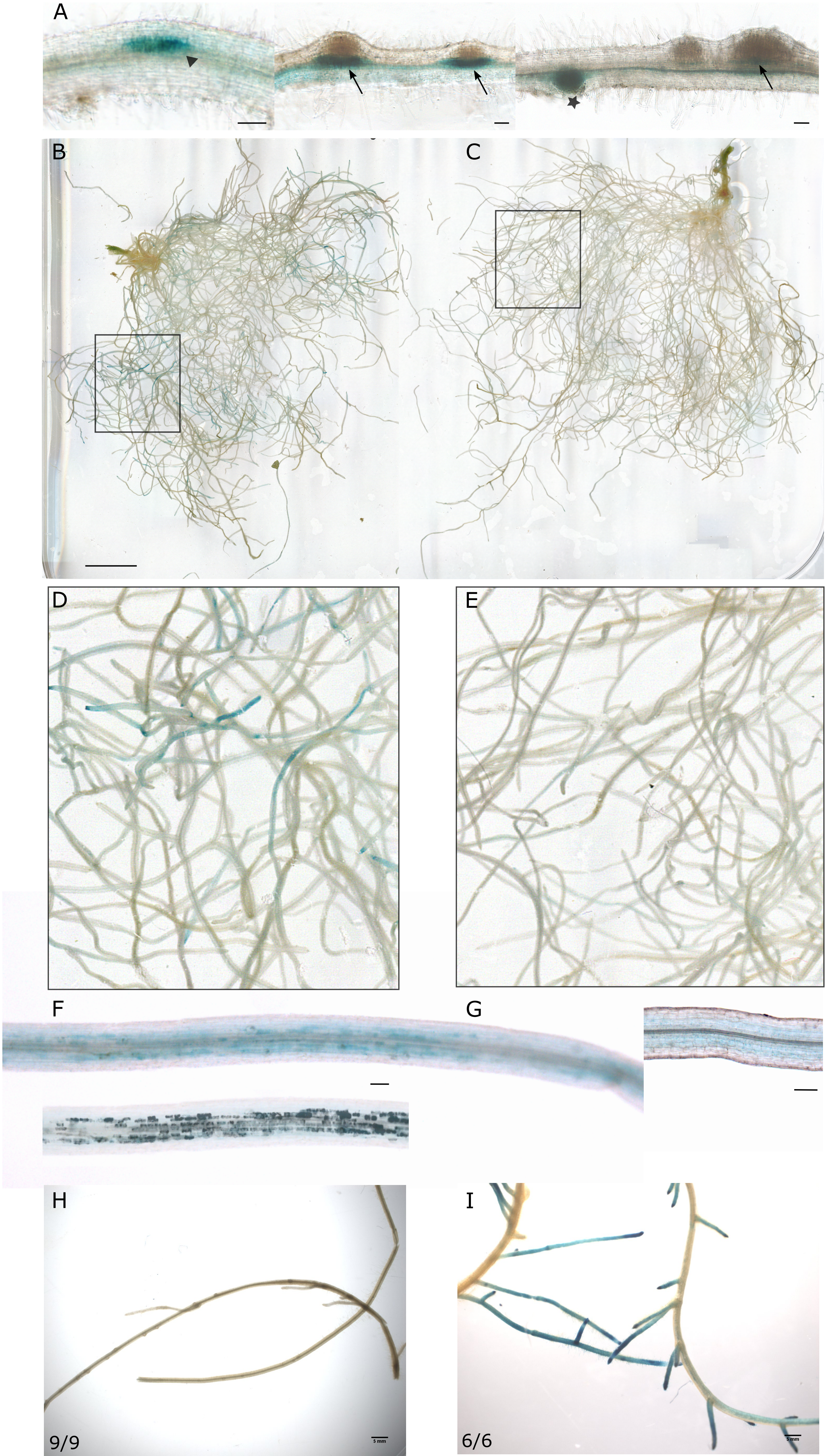

**Supplemental Figure 2: Genomic and proteic alignments of MpaSYMRK in wild type and *M. paleacea* mutant lines.**

**A** and **B.** Genomic alignment of the first and second exon of *MpaSYMRK*. Pairs of sgRNAs used for CRISPR/Cas9 are underlined in different colors. Premature stop codons are underlined in black.
**C.** Proteic alignment of the two first exons of MpaSYMRK from the different mutant lines.
Alignments were performed using Clustal Omega and visualized in AliView.

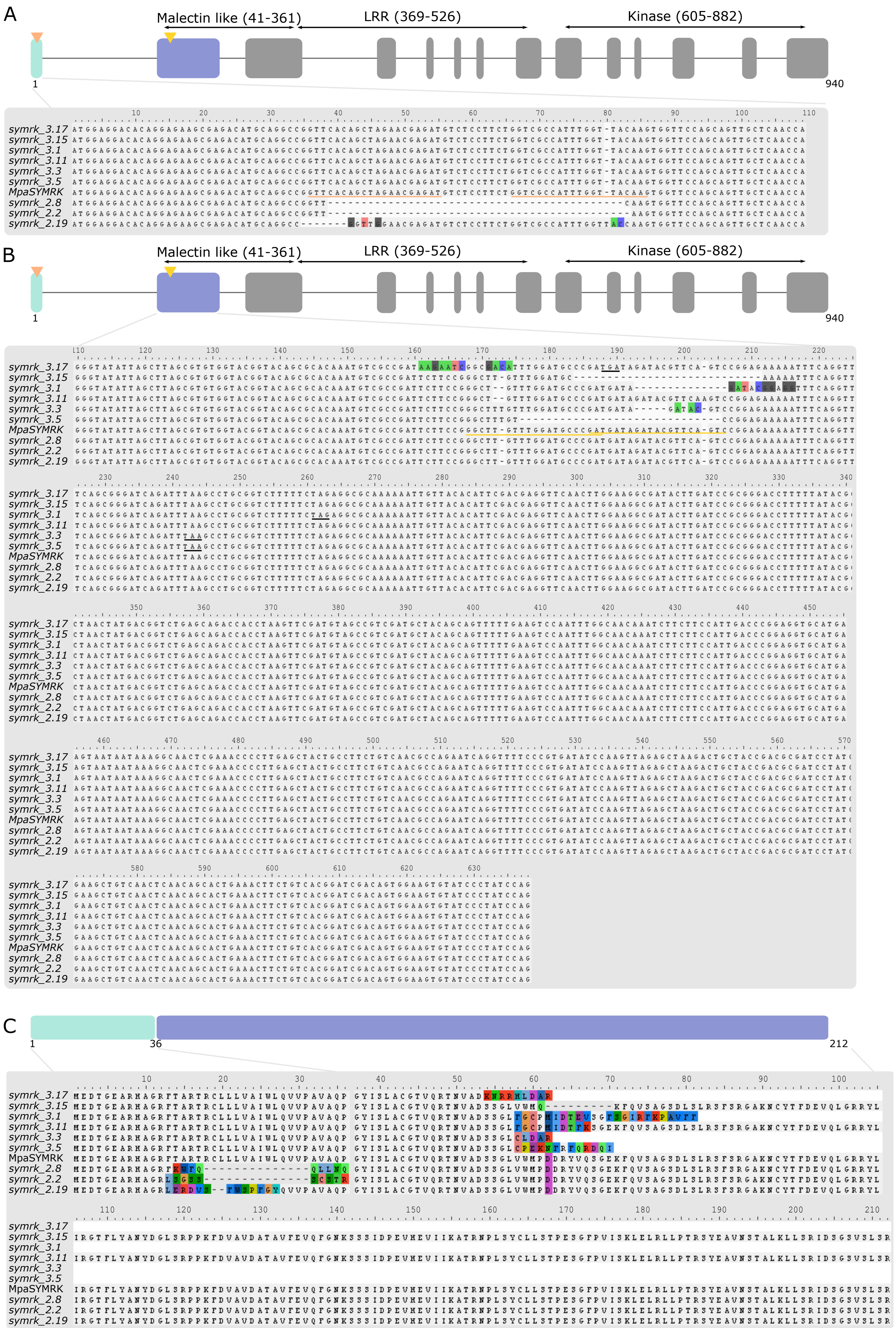

**Supplemental Figure 3: Genomic and proteic alignments of MpaCCaMK in wild type and *M. paleacea* mutant lines.**

**A.** Genomic alignment of the first exon of *MpaCCaMK*. Pairs of sgRNAs used for CRISPR/Cas9 are underlined in different colors. Premature stop codons are underlined in black.

**B.** Proteic alignment of the first exon of MpaCCaMK from the different mutant lines. Alignments were performed using Clustal Omega and visualized in AliView.

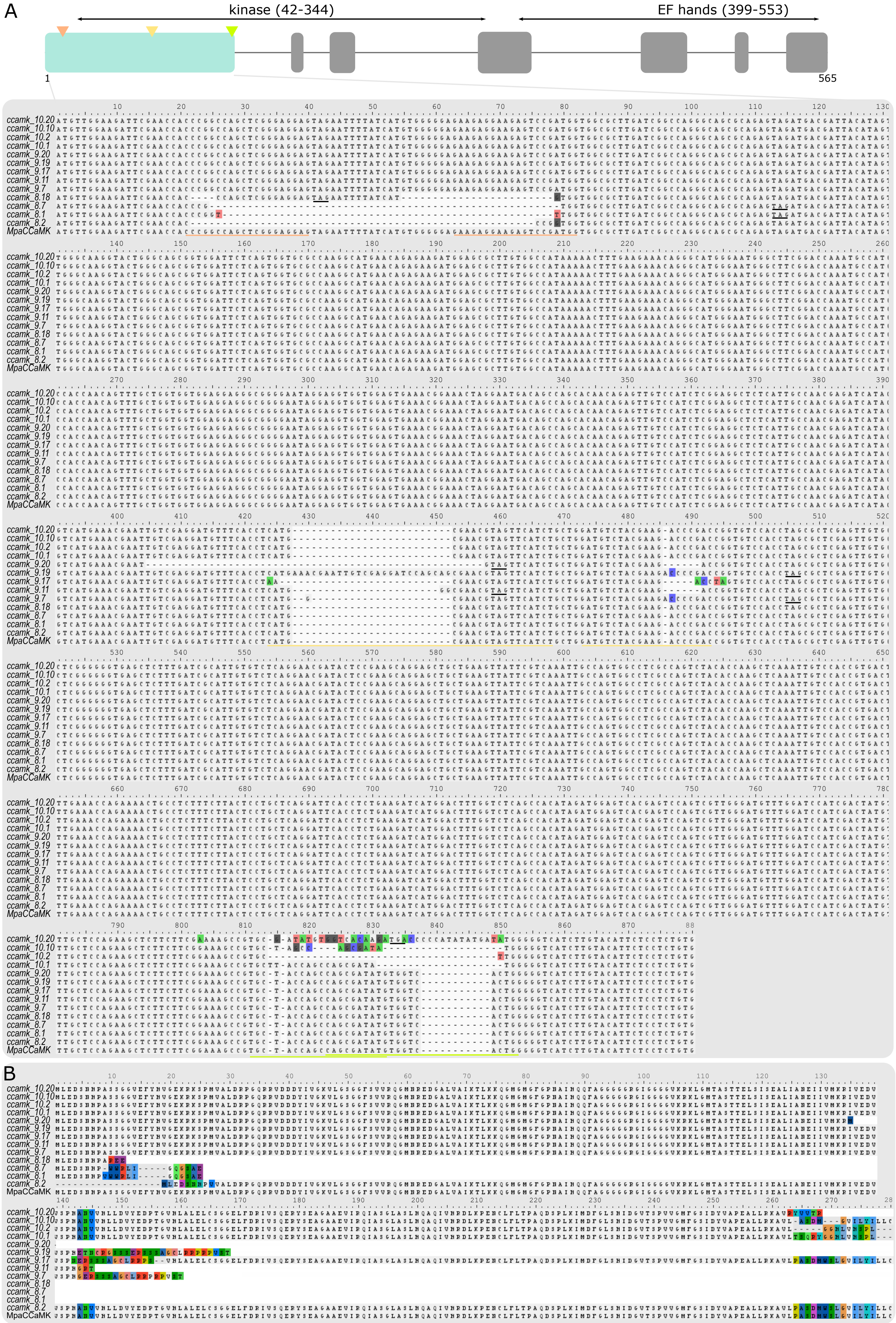

**Supplemental Figure 4: Genomic and proteic alignments of MpaCYCLOPS in wild type and *M. paleacea ssp paleacea* mutant lines.**

**A.** Genomic alignment of the first exon of *MpaCYCLOPS*. Pairs of sgRNAs used for CRISPR/Cas9 are underlined in different colors. Premature stop codons are underlined in black and the predicted new in-frame start codons are underlined in blue. The N-terminal domain absent from angiosperms is indicated in green, the regulatory domain in black.

**B.** Proteic alignment of MpaCYCLOPS first exon until premature stop codon if a stop codon is present.

**C.** Proteic alignment of MpaCYCLOPS first exon from sequences with new predicted in-frame start codons in the different mutant lines. Alignments were performed using Clustal Omega and visualized in AliView.

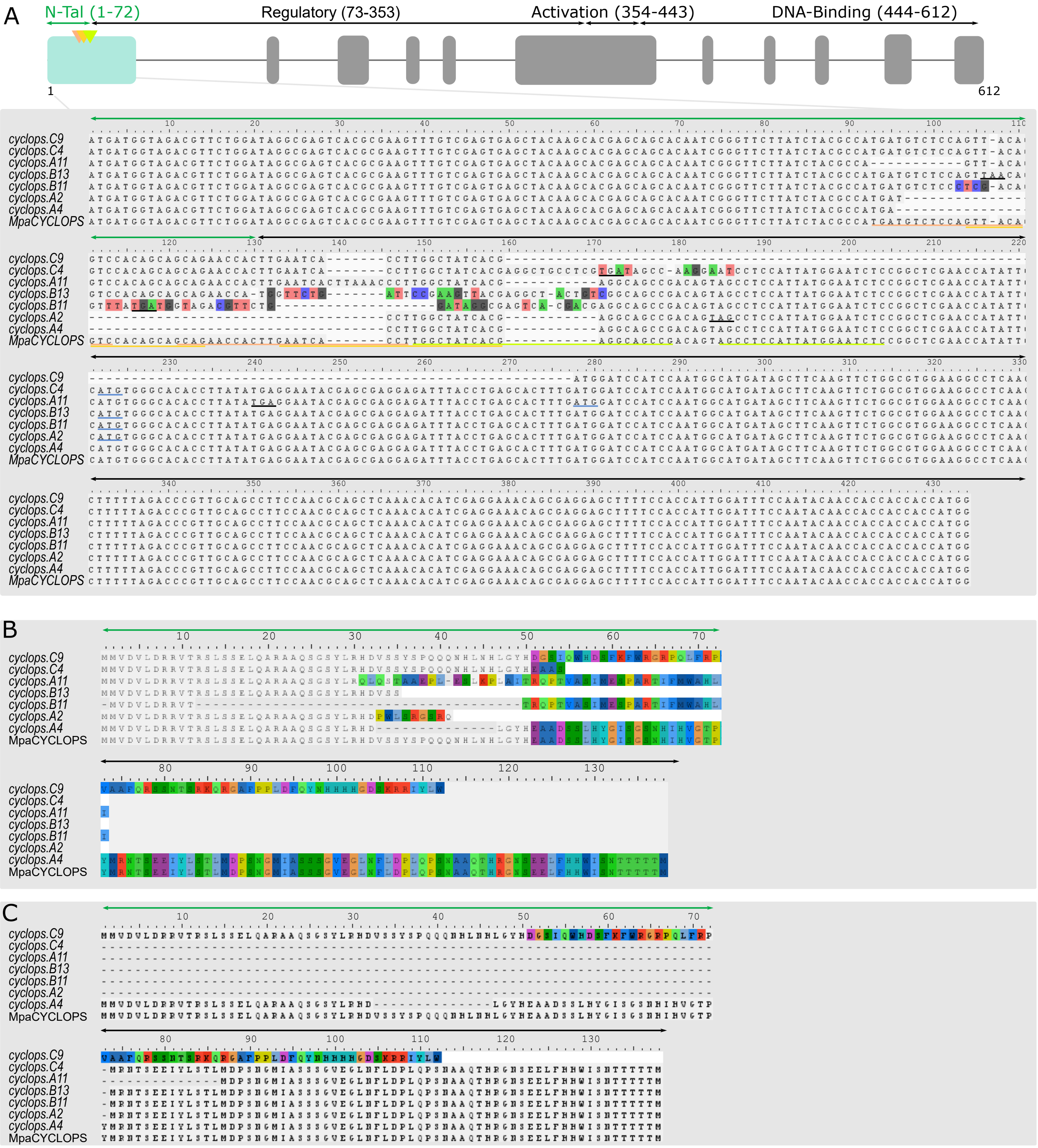

**Supplemental Figure 5: Loss of a N-terminal domain in angiosperms CYCLOPS proteins**

CYCLOPS phylogeny reconstructed using maximum likelihood. The untrimmed alignment of the proteins is shown next to the phylogeny. The N-terminal domain is absent (or highly divergent) in angiosperms and is highlighted in green. The regulatory, activation, and DNA-binding domains are highlighted in red, yellow, and blue, respectively. The two phosphorylation sites are shown with a dark blue rectangle; they were both mutated to D in CYCLOPS-DD overexpressing lines. Ultrafast bootstrap support (%; out of 1,000 replicates) is shown at the end of the branches. The scale represents the estimated number of substitutions per site. Sequences that differ from the original annotation are shown with an asterisk (*), and experimentally validated proteins are marked with a yellow star.

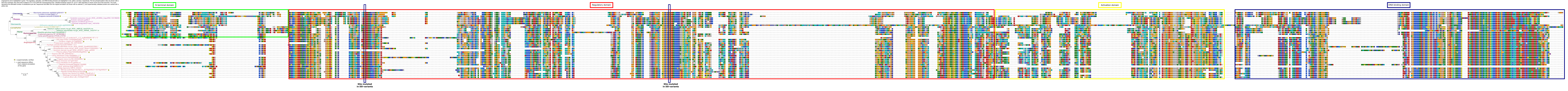

**Supplemental Figure 6:** ***Marchantia paleacea ssp diptera cyclops*** **lines are impaired for arbuscular mycorrhization with** ***R. irregularis***

**A.** Proteic alignments of MpaCYCLOPS first exon in wild type and *M. paleacea ssp paleacea* and *ssp diptera* mutant lines. The N-terminal domain absent from angiosperms is indicated in green, the regulatory domain in black. Alignments were performed using Clustal Omega and visualized in AliView.

**B.** The percentage of plants colonized by *R. irregularis* five weeks post inoculation with *R. irregularis* is indicated. *** indicates statistical difference (p-value<0,001) calculated with a pairwise comparison of proportions (Chi2) to the control line (Control.F) and a BH p-value adjustment. n= number of observed thalli.

**C.** Sections of *cyclops* and control lines five weeks post inoculation with *R. irregularis. R. irregularis* is visualized with blue ink. Scale bar=100µm.

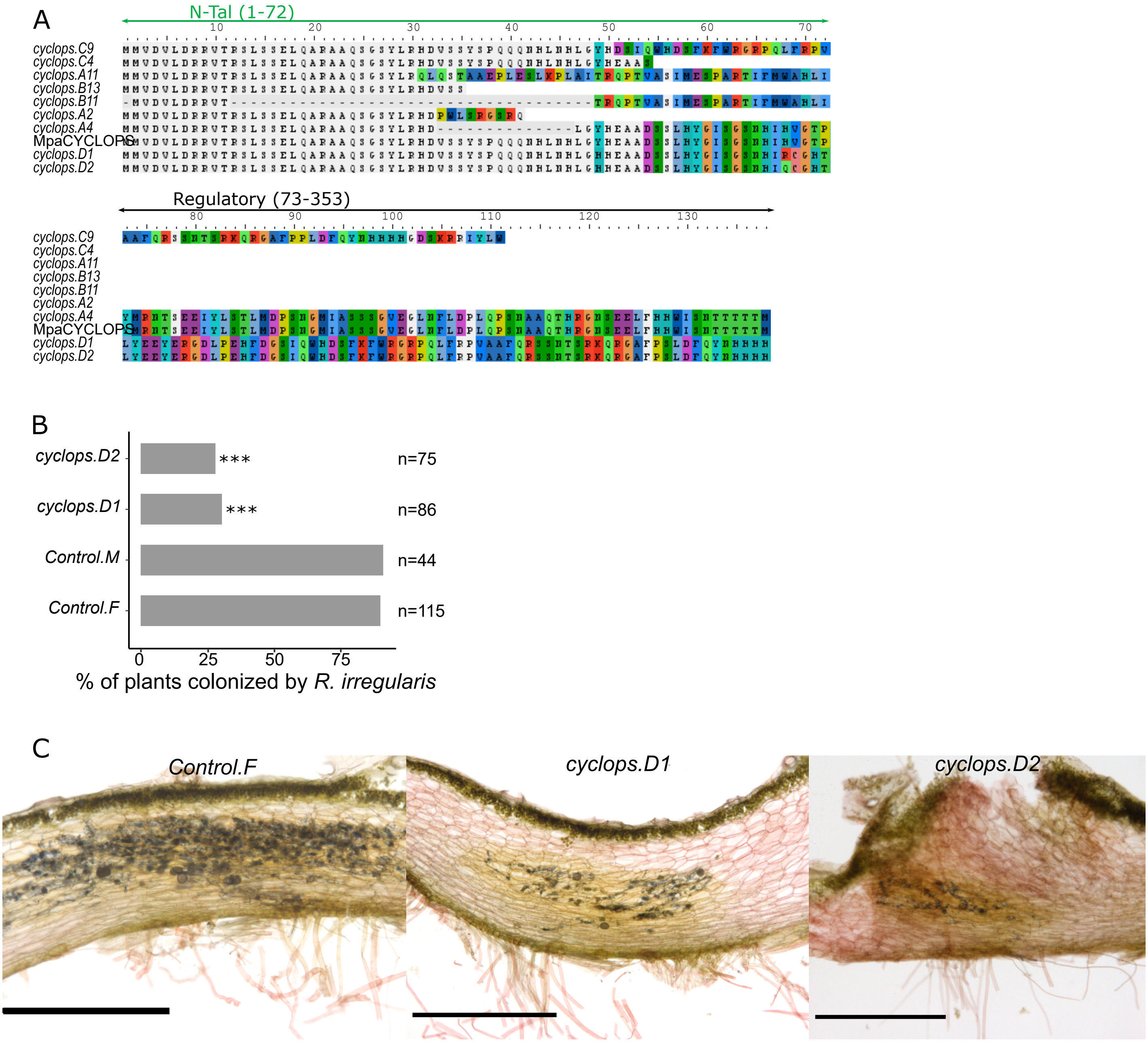
